# Bacterial Modulation of Intestinal Motility through Macrophage Redistribution

**DOI:** 10.1101/2024.05.10.593615

**Authors:** Julia S. Ngo, Piyush Amitabh, Jonah G. Sokoloff, Calvin Trinh, Travis J. Wiles, Karen Guillemin, Raghuveer Parthasarathy

## Abstract

Intestinal microbes, whether resident or transient, influence the physiology of their hosts, altering both the chemical and the physical characteristics of the gut. An example of the latter is the human pathogen *Vibrio cholerae’s* ability to induce strong mechanical contractions, discovered in zebrafish. The underlying mechanism has remained unknown, but the phenomenon requires the actin crosslinking domain (ACD) of *Vibrio’s* Type VI Secretion System (T6SS), a multicomponent protein syringe that pierces adjacent cells and delivers toxins. By using a zebrafish-native *Vibrio* and imaging-based assays of host intestinal mechanics and immune responses, we find that macrophages mediate the connection between the T6SS ACD and intestinal activity: ACD-dependent tissue damage activates macrophages and recruits them from their unperturbed positions near enteric neurons lining the midgut, spurring strong gut contractions resembling those resulting from genetic depletion of macrophages. In addition to illuminating host-directed actions of the widespread T6SS protein apparatus, our findings highlight how localized bacteria-induced injury can reshape neuro-immune cellular dynamics to impact whole-organ physiology.

## Introduction

Intestinal microorganisms, whether commensal or pathogenic, impact their hosts through a wide range of processes (1–6). Though chemical transformations related for example to nutrient metabolism or neurotransmitter synthesis are intensely studied, transformations of the physical environment remain underexplored, likely due to the difficulty of characterizing the gut environment *in situ*. In earlier work (7) we showed that the human pathogen *Vibrio cholerae* (El Tor strain C6706) induces strong mechanical contractions in the intestines of zebrafish, a consequence of the bacterial Type VI Secretion System (T6SS), a syringe-like apparatus with which *Vibrio cholerae*, like nearly 25% of all Gram-negative bacteria, can inject effector proteins into target cells. Enhanced gut contractions, moreover, required the actin crosslinking domain (ACD) present at the C-terminus of the *Vibrio* T6SS spike protein VgrG-1, indicating a eukaryotic target. The nature of this connection between bacterial activity and host mechanical response has, however, remained mysterious.

To map the connection between the T6SS and host intestinal activity, we examined host responses to a *Vibrio* species isolated from and commonly found in zebrafish intestines, strain ZWU0020, which unlike the human-derived *Vibrio cholerae* El Tor C6706, does not encode cholera toxin or toxin-coregulated pilus (8). Cholera toxin can itself induce epithelial barrier damage and diarrhea (9–11). In addition to facilitating a focus on the T6SS, *Vibrio* ZWU0020, hereafter referred to simply as “*Vibrio*” for brevity, colonizes the larval zebrafish gut to about ten times the abundance of *Vibrio cholerae* El Tor C6706 (7), forming dense communities of planktonic, highly motile cells, shown in prior work to be capable of invading existing gut bacterial populations (12) and causing intestinal inflammation in the form of gut-localized neutrophils (13–15) and macrophages expressing the cytokine TNFα (16). The pro-inflammatory character of *Vibrio* contributes to its characterization as a “pathobiont,” a member of the normal microbiota with a potential for pathogenicity (14, 16, 17). We studied commensal *Vibrio* colonizing larval zebrafish, a model whose transparency enables high resolution imaging of the entire intestine including visualization of epithelial tissue and labeled cells, such as immune cells and enteric neurons, in living animals (7, 16, 18–20).

We hypothesized that gut-associated macrophages were a likely intermediary between bacteria and other host cell types because of the above-mentioned observations of *Vibrio*-induced inflammation and because studies using mice and zebrafish have established communication pathways between macrophages and the enteric neurons that regulate muscular activity along the intestine. In mice, Muller *et al*. showed that macrophage secretion of bone morphogenic protein 2 (BMP2) activates enteric neurons, leads to hyperactive contractions in intestinal segments *ex vivo*, and increases colonic transit time (21). In zebrafish, Graves *et al*. found intestinal macrophages in close proximity to enteric neurons and showed that genetic ablation of the *irf8* gene, previously shown to be necessary for macrophage differentiation (22), leads to shorter transit times in larvae, though not in adults (23). How bacterial activity, especially mediated by the T6SS, might intersect with macrophage activity or mortality has remained unclear.

The imaging-based assays described here, performed in transgenic animals that allow manipulation and visualization of macrophages, revealed host tissue damage, inflammation, and changes in gut contraction dynamics induced by the *Vibrio* T6SS ACD. Strikingly, we found an ACD-dependent spatial reorganization of macrophages that appears to drive strong gut contractions while leaving the frequency of contractions unchanged. These findings not only provide insights into the ways by which resident microbes influence their hosts but also indicate an unexpected and potentially general mechanism for the regulation of intestinal mechanics via physical reorganization of relevant cell types.

## Results

### The *Vibrio* T6SS actin crosslinking domain is required for enhanced intestinal contractions

To determine the role of the bacterial T6SS, specifically the eukaryote-targeting actin crosslinking domain (ACD), we generated a mutant of the zebrafish-native *Vibrio* with an in-frame deletion of the ACD of the T6SS spike protein VgrG-1 (Methods); we will refer to this strain as *Vibrio*^ΔACD^. We also generated a mutant strain lacking the entire T6SS gene cluster, denoted *Vibrio*^ΔT6SS^ (Methods). The wild-type *Vibrio* and both mutants show identical *in vitro* growth rates in lysogeny broth (Figure S1).

We confirmed that deletion of the ACD does not interfere with the functionality of the Type VI apparatus, other than through effects directly mediated by the ACD itself, by checking that *Vibrio*^ΔACD^, but not *Vibrio*^ΔT6SS^, is capable of killing bacterial targets. We performed an *in vitro* assay in which one of the *Vibrio* strains was spotted onto agar plates along with another zebrafish-native species, the previously examined *Aeromonas* ZOR0001 (7, 12) (Methods). When co-spotted with the completely T6SS-deficient *Vibrio*^ΔT6SS^, *Aeromonas* ZOR0001 is present at high abundances that increase monotonically with the concentration of the initial suspension, as when spotted alone, consistent with a lack of inter-bacterial killing (Figure S2). In contrast, *Aeromonas* ZOR0001 co-spotted with *Vibrio*^ΔACD^ shows consistently low abundances that drop to nearly zero at high initial concentrations, as when spotted with wild-type *Vibrio*, consistent with inter-bacterial killing (Figure S2).

*In vivo*, both the wild-type *Vibrio* and *Vibrio*^ΔACD^, when inoculated in mono-association with initially germ-free larval zebrafish, colonized the gut to approximately the same abundance, with the mean ± standard deviation of log_10_(bacteria per gut) being 5.0 ± 0.4 and 5.2 ± 0.4 for wild-type *Vibrio* and *Vibrio*^ΔACD^, respectively (Figure S3).

Larval zebrafish, like other animals, exhibit periodic intestinal contractions that propagate along the length of the gut. As in past work, imaging using differential interference contrast (DIC) microscopy and analysis based on image velocimetry reveal and quantify the frequency and strength of these contractions (7, 20). Germ-free (GF) fish were inoculated with either *Vibrio* or *Vibrio*^ΔACD^ at 5 days post-fertilization (dpf). The next day, at 24-30 hours post-inoculation (hpi), we used DIC microscopy to acquire movies of 5 minute duration at 5 frames per second spanning a roughly 400 μm segment of the posterior intestine of live fish (7, 20), indicated in Figure 1A. It is readily visually apparent that contractions are stronger in fish inoculated with wild-type *Vibrio* compared to *Vibrio*^ΔACD^ (Movies S1, S2). To quantify intestinal mechanics, we performed image velocimetry to determine the velocity field at a grid of points spanning the images (Figure 1B, C). The cross-correlation of the velocity field shows periodic, propagating waves with a frequency revealed by Fourier analysis. The amplitude of the image velocity at the dominant frequency provides a measure of the strength of the contractions, essentially the amplitude of tissue motion corresponding to rhythmic oscillations (20). Zebrafish mono-associated with wild-type *Vibrio* showed on average roughly 100% larger contraction amplitude as those mono-associated with *Vibrio*^ΔACD^ (Figure 1D; ratio 1.9 ± 0.6, p = 0.039 from a non-parametric Mann-Whitney U test), similar to effects seen previously with human-derived *Vibrio cholerae* El Tor C6706 (7). Notably, the frequency of contractions was very similar for the two bacterial conditions (Figure 1E), being 1.92 ± 0.04 1/min. (mean ± s.e.m.) for wild-type *Vibrio* and 2.04 ± 0.09 1/min. for *Vibrio*^ΔACD^ (p = 0.63 from a non-parametric Mann-Whitney U test).

**Figure 1.**
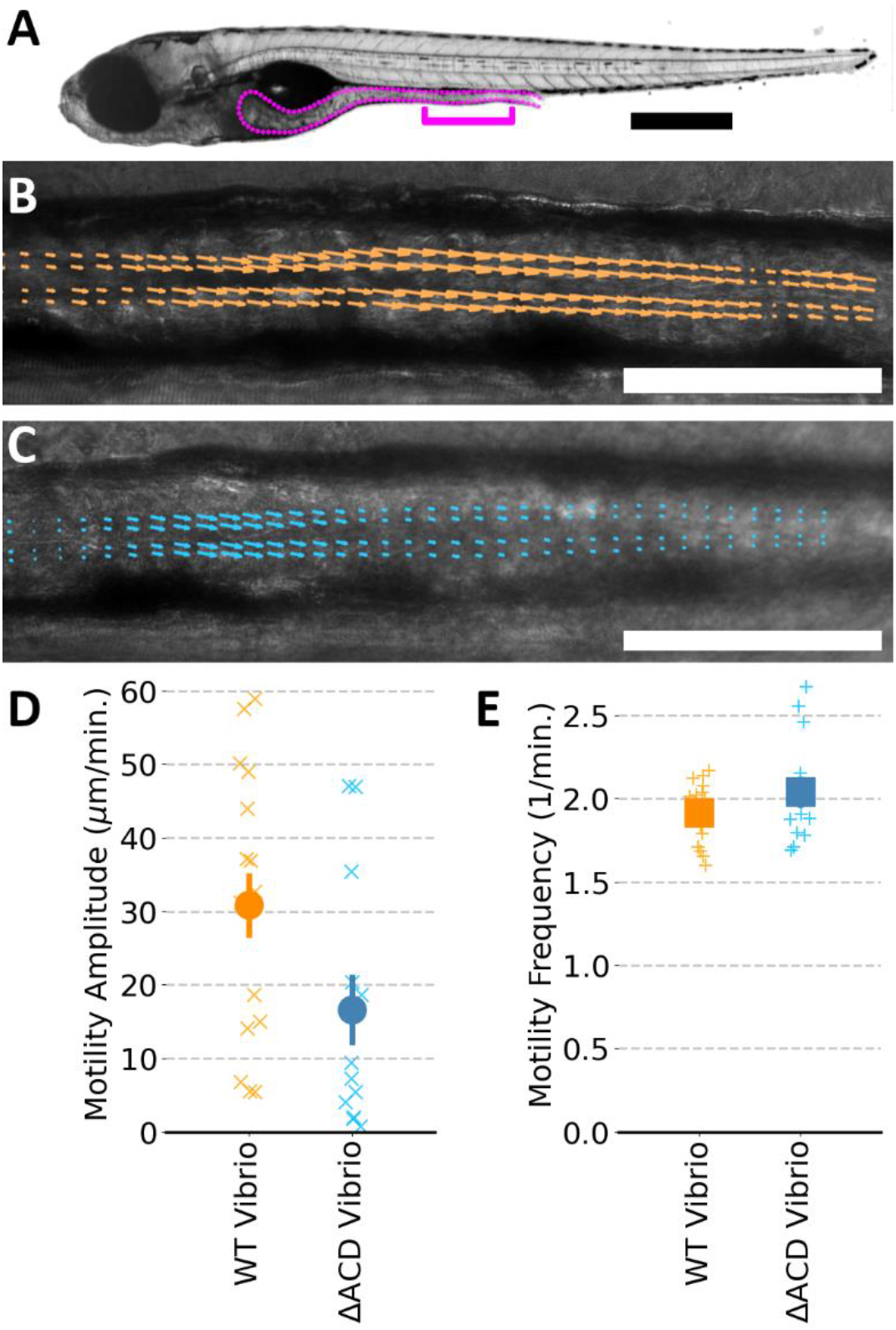
(A) Brightfield image of a larval zebrafish at 6 dpf, with the gut outlined. The bracket indicates the region imaged for intestinal motility assessment. Bar: 500 μm. (B, C) DIC images of a 6 dpf zebrafish, initially germ-free and inoculated at 5 dpf with (B) wild-type *Vibrio* and (C) *Vibrio*^ΔACD^ with superimposed arrows from image velocimetry analysis. In both images, the magnitude of arrows is assigned using the same velocity scale, the selected zebrafish are near the median motility amplitude for their condition, and the selected timepoint is near the peak of an intestinal contraction. Bar: 100 μm. See also movies S1 and S2. (D) Gut motility amplitudes of 6 dpf zebrafish. The amplitude of periodic intestinal contractions is nearly 100% larger (ratio 1.9 ± 0.6) for fish mono-associated with wild-type *Vibrio* compared to *Vibrio*^ΔACD^ (p = 0.039 from a non-parametric Mann-Whitney U test). (E) The mean frequency of intestinal contractions is very similar between the groups (mean ± s.e.m. = 1.92 ± 0.04 1/min. for wild-type *Vibrio*, 2.04 ± 0.09 for *Vibrio*^ΔACD^; p = 0.63). In (D) and (E), “x”s indicate measurements of individual zebrafish; solid symbols and error bars indicate the mean and standard error of the mean, respectively.

### Macrophages downregulate gut motility and are necessary for ACD-dependent enhancement of gut contractions

We suspected macrophages as an intermediary between *Vibrio* activity and host intestinal motility. To quantitatively characterize the impact of macrophages on propagating intestinal contractions, we generated zebrafish with depleted numbers of macrophages. In brief, we used CRISPR-Cas9 to target the *irf8* gene, crucial for the differentiation of macrophages (23), injecting embryos at the single-cell stage with the Cas9 enzyme and the appropriate single guide RNAs to yield F0 “crispants” (24), which we will denote Δ*irf8* (Methods). Use of a transgenic zebrafish line with fluorescently labeled macrophages (*Tg(mpeg1:mCherry)*) allows quantification of macrophage number. We find that the Δ*irf8* zebrafish at 6 dpf have roughly half as many macrophages as otherwise wild-type fish (Figure 2A-C).

**Figure 2.**
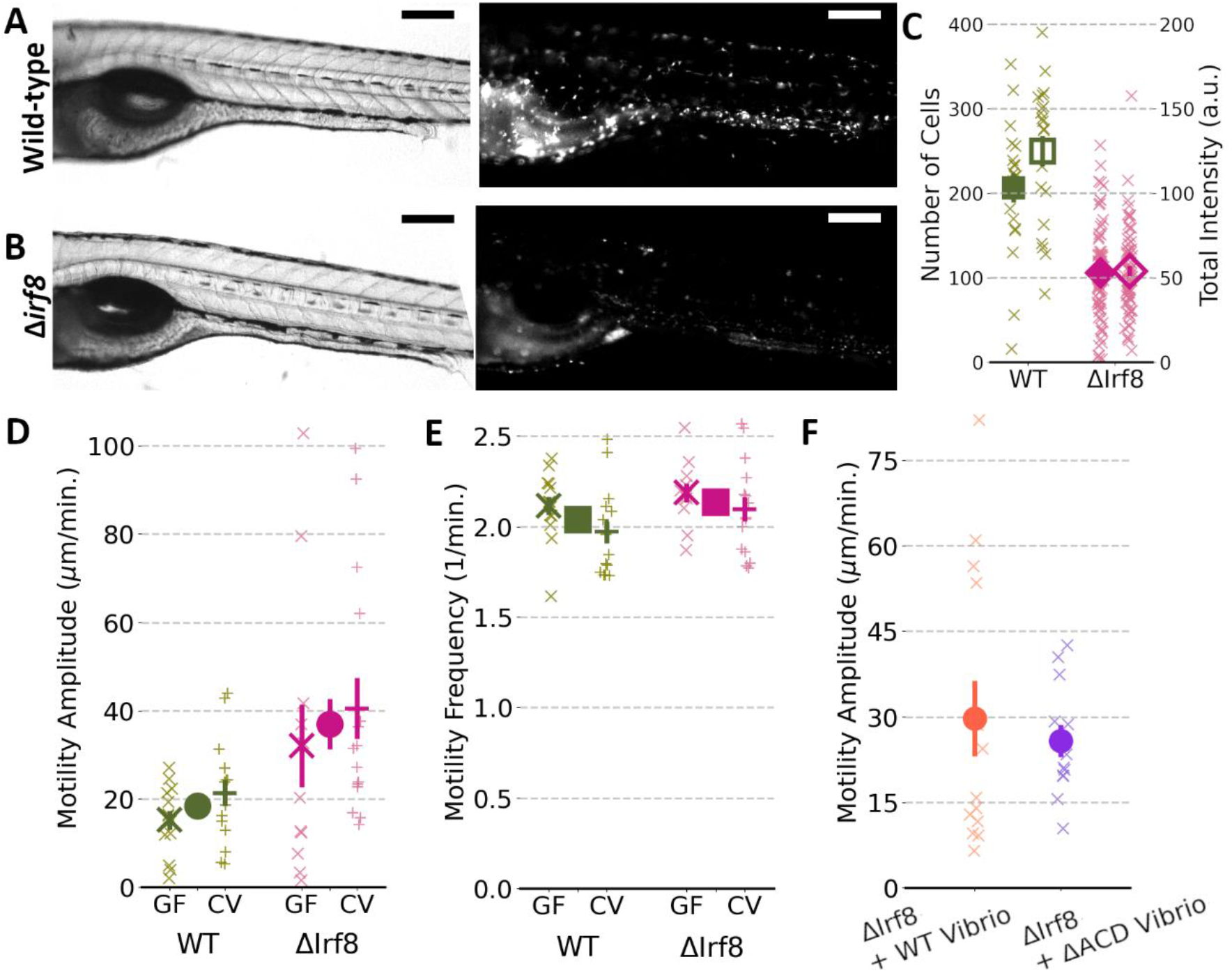
(A) Brightfield (left) and fluorescence (right) images of the mid-section of a 6 dpf (*Tg(mpeg1:mCherry)*) zebrafish with fluorescent macrophages. Bar = 200 μm. (B) Brightfield and fluorescence images from a 6dpf Δ*irf8* zebrafish. The fluorescence image is shown at the same intensity scale as in (A). Bar = 200 μm. (C) Quantification of macrophage abundance by number of fluorescent cells (solid symbols) and total background-subtracted mCherry fluorescence (open symbols). Each “x” represents a measurement from an individual zebrafish over the mid-section as in (A, B). Each large symbol represents the mean, with error bars indicating the standard error of the mean (s.e.m.). The fish shown in (A, B) were selected for proximity of their macrophage counts to the median value of the population. (D) Gut motility amplitudes of 6 dpf wild-type and Δ*irf8* zebrafish. Each small “x” or “+” represents a measurement from an individual germ-free (GF) or conventionally reared (CV) zebrafish, respectively, with the large “x” or “+” indicating the mean ± s.e.m. The central circles and error bars for each fish type denote the mean and s.e.m. of the pooled data from germ-free and conventional fish, which show the amplitude of periodic intestinal contractions to be 100% larger (ratio 2.0 ± 0.4) for fish with depleted macrophages (Δ*irf8*) compared to wild-type fish (p = 0.016 from a non-parametric Mann-Whitney U test). (E) The mean frequency of intestinal contractions is approximately equal for the two groups (mean ± s.e.m. = 2.04 ± 0.04 for wild-type zebrafish, 2.13 ± 0.04 for Δ*irf8* zebrafish, similarly pooled as in (D); p = 0.13). (F) Gut motility amplitudes of initially germ-free 6 dpf Δ*irf8* zebrafish mono-associated with wild-type *Vibrio* or *Vibrio*^ΔACD^ are similar to each other (ratio 1.2 ± 0.4; p = 0.58 from a non-parametric Mann-Whitney U test). As in previous panels, “x”s indicate measurements of individual zebrafish; solid symbols and error bars indicate the mean and standard error of the mean, respectively.

It has been shown that larval zebrafish lacking the *irf8* gene have decreased intestinal transit time for solid food compared to wild-type siblings (23). However, the relationship between transit time and intestinal mechanics is not straightforward, as it may depend on the frequency of contractions, their strength, their coherence, and other factors. We imaged and analyzed intestinal contractions in 6 dpf wild-type or Δ*irf8* fish. Whether germ-free or conventionally reared, the macrophage-depleted Δ*irf8* fish showed highly variable contraction amplitudes with a mean roughly twice as large as that of wild-type fish (Figure 2D). The ratio of amplitudes of Δ*irf8* / wild-type fish is 2.1 ± 0.7 and 1.9 ± 0.4 for germ-free (GF) and conventional (CV) fish, respectively (Figure 2D). Pooling the GF and CV data, the ratio of gut contraction amplitudes of Δ*irf8* / wild-type fish overall is 2.0 ± 0.4; p = 0.016 from a non-parametric Mann-Whitney U test (Figure 2D). In contrast to the amplitude, the frequency of contractions was nearly identical for the two types of fish (Figure 2E), being 2.04 ± 0.04 1/min. (mean ± s.e.m.) for wild-type fish and 2.13 ± 0.04 1/min. for Δ*irf8* fish (pooled GF and CV; p = 0.13), matching the above-mentioned values from wild-type fish mono-associated with *Vibrio*.

To test whether a normal macrophage abundance is necessary for ACD-mediated enhancement of intestinal motility, Δ*irf8* fish were derived germ-free and then mono-associated with either wild-type *Vibrio* or *Vibrio*^ΔACD^. As above, intestinal dynamics were assessed the day after inoculation. Contraction amplitudes were similar for both sets (Figure 2F; ratio 1.2 ± 0.4; p = 0.58) and similar to the value measured for Δ*irf8* fish in the absence of *Vibrio*, implying that macrophages and the *Vibrio* ACD are part of the same pathway for increasing intestinal contractions strength. Again, contraction frequencies were independent of condition (2.05 ± 0.08 1/min. (mean ± s.e.m.) for wild-type fish and 1.98 ± 0.04 1/min. for Δ*irf8* fish).

### The *Vibrio* T6SS actin crosslinking domain is required for macrophage activation

Prior work demonstrated that *Vibrio* stimulates the innate immune system as reported by production of the proinflammatory cytokine TNFα (16). To clarify the role of the T6SS ACD, we derived germ-free transgenic zebrafish with fluorescent reporters of macrophages and TNFα expression (*Tg(TNF*α:*GFP);Tg(mpeg1:mCherry)*), inoculated the fish with bacteria, and dissected their intestines at 24 hours post-inoculation to focus on gut-associated signals (Figure 3A). In addition to broad epithelial TNFα:GFP fluorescence, discrete GFP-positive cells were evident (Figure 3A), the number of which was nearly three times greater in fish inoculated with wild-type *Vibrio* compared to *Vibrio*^ΔACD^ or to germ-free fish (Figure 3B). In detail: the mean ± s.e.m. of GFP+ cells was 18.1 ± 1.9 for fish with wild-type *Vibrio* (*N* = 13), 7.3 ± 2.3 for fish with wild-type *Vibrio*^ΔACD^ (*N* = 16), and 5.3 ± 1.7 for germ-free fish (*N* = 9); comparing wild-type *Vibrio* and *Vibrio*^ΔACD^ gives p = 0.002 from a non-parametric Mann-Whitney U test. Moreover, the number of GFP-positive and mCherry-positive cells, i.e. macrophages expressing *TNF*α, was six times greater in fish inoculated with wild-type *Vibrio* compared to *Vibrio*^ΔACD^ or to germ-free fish (Figure 3C). In detail: the mean ± s.e.m. of GFP+ and mCherry+ cells was 7.8 ± 1.2 for fish with wild-type *Vibrio* (*N* = 13), 1.3 ± 0.4 for fish with wild-type *Vibrio*^ΔACD^ (*N* = 16), and 1.2 ± 0.4 for germ-free fish (*N* = 9); comparing wild-type *Vibrio* and *Vibrio*^ΔACD^ gives p = 0.0002 from a non-parametric Mann-Whitney U test. The ACD, therefore, is crucial for macrophage activation.

**Figure 3.**
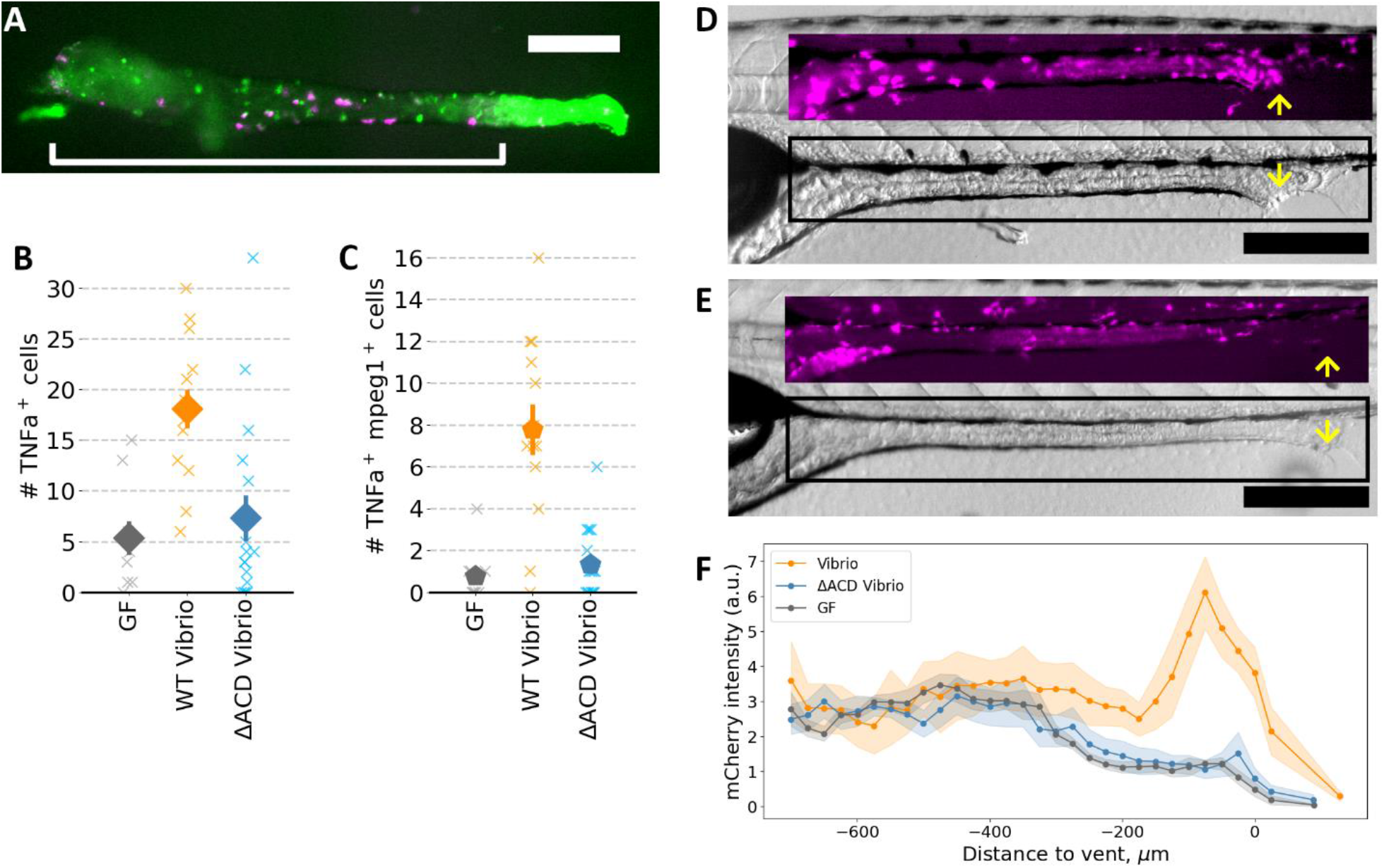
(A) Representative image of a dissected gut from a 6 dpf Tg*(tnfa*:GFP);Tg*(mpeg1*:mCherry) zebrafish mono-associated with wild-type *Vibrio*, showing gut-associated macrophages (mCherry^+^) and cells expressing the *tnfa* reporter (GFP^+^). The bracket indicates the region assessed for cell counts. Bar: 250 μm. (B) Numbers of gut-associated *tnfa*-positive cells in germ-free fish, fish mono-associated with wild-type *Vibrio*, and fish mono-associated with *Vibrio*^ΔACD^. (C) Numbers of gut-associated *tnfa*-positive macrophages (*tnfa*:GFP^+^ and *mpeg1*:mCherry^+^). In (B) and (C), “x”s indicate measurements of individual zebrafish; solid symbols and error bars indicate the mean and standard error of the mean, respectively. (D, E) Representative brightfield and macrophage-fluorescence (insets) images of 6 dpf Tg*(tnfa*:GFP);Tg*(mpeg1*:mCherry) zebrafish mono-associated with (D) wild-type *Vibrio* and (E) *Vibrio*^ΔACD^. The rectangle indicates the inset region. Yellow arrows indicate the vent. Bar: 250 μm. (F) Mean mCherry intensity averaged over the gut as a function of anterior-posterior position, relative to the vent, for germ-free (GF) zebrafish (*N*=6), zebrafish mono-associated with wild-type *Vibrio* (*N*=6), and zebrafish mono-associated with *Vibrio*^ΔACD^ (*N*=8). Shaded bands indicate the standard error of the mean at each position.

We also noticed through simple epifluorescence imaging high mCherry (macrophage) intensity near the zebrafish posterior vent (the posterior opening of the gut) for fish mono-associated with wild-type *Vibrio* and not for other conditions (Figure 3D-F), motivating closer examination of tissue phenotypes and cellular organization in this region.

### *Vibrio* causes posterior tissue damage but does not preferentially kill macrophages

Comparing at 6 dpf initially germ-free zebrafish inoculated 24 hours prior with *Vibrio* or *Vibrio*^ΔACD^, or kept germ-free, we observed a pronounced enlargement of the vent (Figure 4A), which was on average 2.5 times wider in fish colonized with *Vibrio* (Figure 4B). Live imaging starting at 13 hpi showed steady, progressive vent widening and the increased occurrence of rounded, disordered cells in *Vibrio*-inoculated fish (Figure 4 B, C).

**Figure 4.**
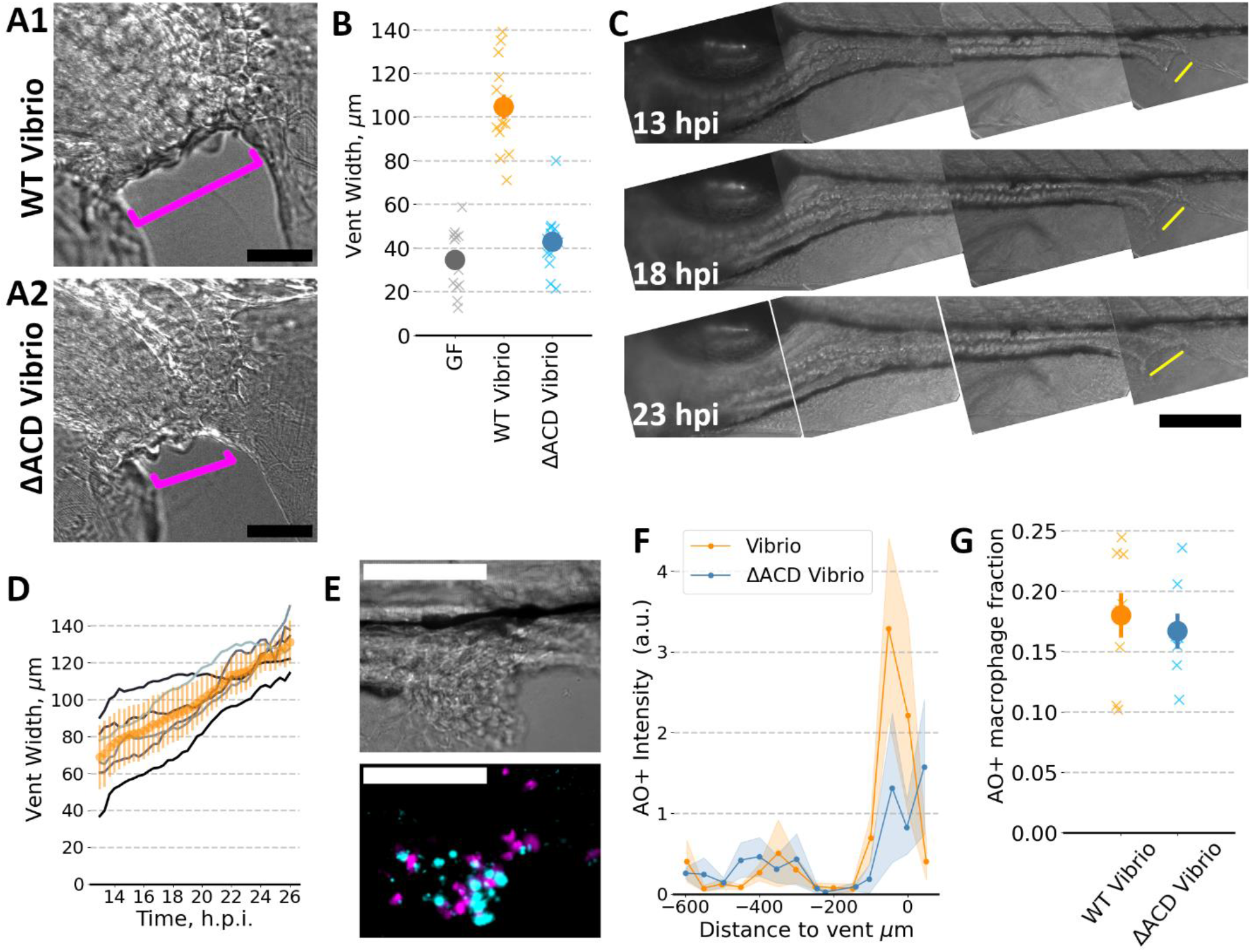
(A) Representative bright-field images of the vent region of 6 dpf zebrafish mono-associated with wild-type *Vibrio* or with ΔACD-*Vibrio*, both at 24 hours post-inoculation. Bar: 50 μm. (B) Vent width of 6 dpf zebrafish, germ-free or mono-associated with wild-type *Vibrio* or with *Vibrio*^ΔACD^, the latter two measured at 24 hpi. For wild-type *Vibrio* compared to *Vibrio*^ΔACD^, the ratio of mean vent widths is larger by a factor of 2.4 ± 0.3 (p = 3.2 × 10^−5^ from a non-parametric Mann-Whitney U test). “X”s indicate measurements of individual zebrafish; solid symbols and error bars indicate the mean and standard error of the mean, respectively. (C) Brightfield images of the gut of a larval zebrafish mono-associated with wild-type *Vibrio* at 13, 18, and 23 hpi (top to bottom). Yellow lines, offset from the vent, indicate the vent width. Bar: 50 μm. (D) Vent width over time for six zebrafish mono-associated with wild-type *Vibrio*. Gray: data from individual fish; Orange: mean, with error bar indicating the standard deviation. (E) Bright field (upper) and fluorescence (lower) images of the vent region of a representative zebrafish mono-associated with wild-type *Vibrio*. Magenta (*mpeg1*:mCherry) indicates macrophages, and cyan (acridine orange) preferentially labels apoptotic cells. Bar: 100 μm. (F) The total fluorescence intensity of all acridine orange-positive cells, binned by anterior-posterior position relative to the vent, for zebrafish mono-associated with wild-type *Vibrio* or with *Vibrio*^ΔACD^. Solid lines indicate average values and shaded bands show the standard error of the mean for *N* = 8 and *N* = 7 fish for wild-type *Vibrio* or with *Vibrio*^ΔACD^, respectively. (G) The fraction of macrophages that are acridine orange-positive for zebrafish mono-associated with wild-type *Vibrio* or with *Vibrio*^ΔACD^. “X”s indicate measurements of individual zebrafish; solid symbols and error bars indicate the mean and standard error of the mean, respectively; p = 0.79 from a non-parametric Mann-Whitney U test.

Suspecting host cell death, we stained live fish with acridine orange (Methods), a membrane-permeable dye that fluoresces in acidic lysosomal vesicles, thereby preferentially staining apoptotic cells (25). Examination of the vent revealed considerable acridine orange signal in fish colonized with *Vibrio* compared to *Vibrio*^ΔACD^ (Figure 4E, F), indicating ACD-mediated cell death.

We hypothesized that macrophages may be prevalent among the dead cells, both because of the demonstrated ability of bacteria to kill macrophages using the T6SS ACD in *in vitro* cultures (26) and because such an effect would reduce the number of macrophages and therefore, like in the Δ*irf8* fish, could explain increased gut contractions. However, examining the fraction of macrophages (*mpeg1:mCherry*+ cells) that showed acridine orange staining, we found no difference between fish colonized with *Vibrio* compared to *Vibrio*^ΔACD^, the mean fraction being less than 20% in either case (Figure 4G). *Vibrio*-mediated cell killing via the ACD therefore does not specifically deplete macrophages.

### *Vibrio* induces redistribution of macrophages

Though simple assessment of mCherry fluorescence intensity in *mpeg1:mCherry* fish suggested that *Vibrio* induced greater macrophage numbers at the vent (Figure 3 D-F), epifluorescence imaging cannot resolve individual cells and so cannot distinguish between greater numbers and greater *mpeg1* expression. We used light sheet fluorescence microscopy to image 6 dpf larval zebrafish all fixed in paraformaldehyde at 24 hours post-inoculation, segmenting the resulting three-dimensional image stacks to identify each macrophage (Methods). Fish inoculated with *Vibrio*, compared to *Vibrio*^ΔACD^ or germ-free fish, showed a strong peak in macrophage density in the vicinity of the vent, over two times greater than the other conditions (Figure 5A).

**Figure 5.**
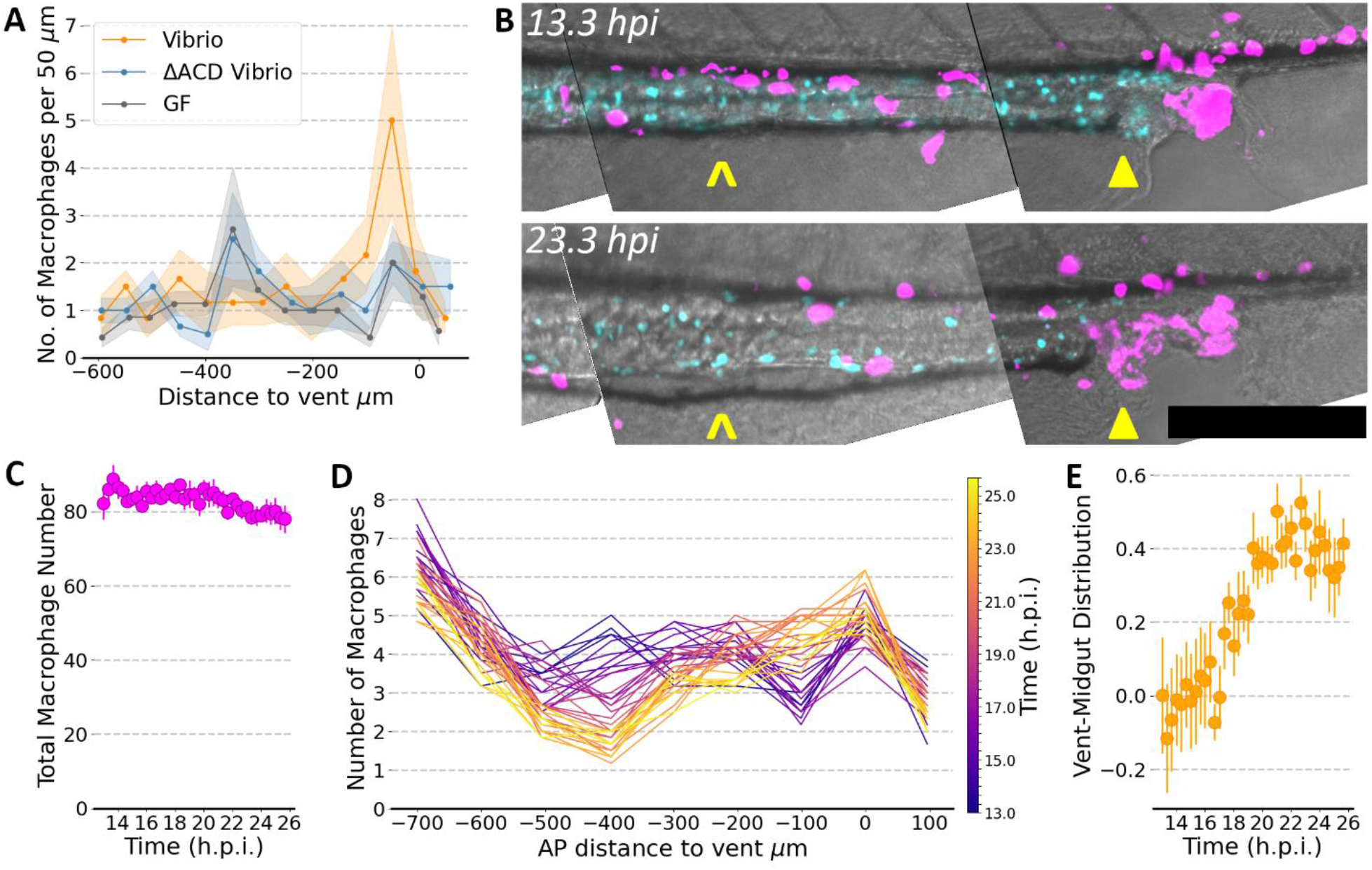
(A) The spatial distribution of macrophages in larval zebrafish, germ-free (GF) or mono-associated with wild-type *Vibrio* or with *Vibrio*^ΔACD^, binned by anterior-posterior position relative to the vent. Fish were fixed and imaged 24 hours post-inoculation, or at the equivalent time for germ-free fish. Solid lines indicate average values and shaded bands show the standard error of the mean for *N* = 6 and *N* = 6 fish for wild-type *Vibrio* or with *Vibrio*^ΔACD^, respectively, and *N* = 7 germ-free fish. (B) Representative composite images from live imaging of larval zebrafish mono-associated with wild-type *Vibrio*. The images shown, maximum intensity projections of the posterior 800 μm of the gut at 13.3 hpi (upper) and 22.3 hpi (lower), are subsets of datasets that comprise three-dimensional image stacks that span the entire gut, acquired every 20 minutes for 13 hours. Magenta (*mpeg1*:mCherry) indicates macrophages, and cyan (*phox2bb*:GFP) indicates enteric neurons; gray is brightfield. Note the depletion of macrophages in the midgut (carat symbols) and the accumulation near the vent (solid triangles). Bar: 200 μm. (C-E) Quantification of cell numbers and positions assessed from identification of cells in volumes spanning the entire gut from live imaging of *N* = 6 fish mono-associated wild-type *Vibrio*. (C) The total number of macrophages over time; circles indicate the mean; bars indicate the standard error of the mean. (D) The spatial distribution of macrophages over time. Each curve indicates the number of macrophages, averaged over *N* = 6 fish, as a function of anterior-posterior position relative to the vent, with color indicating time from 13 to 26 hpi. (E) The relative distribution of macrophages as a function of time, shown as the number of macrophages in a 200 μm posterior region (−150 to +50 μm relative to the vent) minus the number in a 200 μm midgut region (−500 to -300 μm), normalized by the total number of macrophages in these regions.

The above observations spurred us to monitor macrophage positions in live fish over the entire larval gut and vicinity. The rapid acquisition speed of light sheet fluorescence microscopy allows imaging without blurring from peristaltic motions, and its low phototoxicity enables imaging over long durations (18, 27–29). We imaged 5 dpf transgenic fish with labeled macrophages and enteric neurons (*Tg(mpeg:mCherry);Tg(phox2bb:GFP)* (30, 31)), mono-associated with *Vibrio*, from 13 hpi to 26 hpi at intervals of 20 minutes. We observed accumulation of macrophages near the vent as time progressed. Strikingly, this accumulation was accompanied by a depletion of macrophages from the midgut, leaving the enteric neurons lining the gut in this region relatively devoid of nearby macrophages (Figure 5B). Quantifying the macrophage numbers and spatial distribution verified that the total number of macrophages remained roughly constant over the imaging duration (Figure 5C), while the midgut population steadily declined as the posterior vent population increased (Figure 5D, E).

## Discussion

Through a series of experiments based on imaging and genetic manipulation of both bacteria and host, we examined the link between a dramatic amplification of intestinal mechanical activity and a eukaryote-targeting bacterial effector, the Type VI Secretion System actin crosslinking domain. We observed that depletion of macrophages and the presence of the ACD each spur a roughly 100% increase in gut contraction strength that is not further enhanced by imposing both factors together. In contrast to their impact on the magnitude of contractions, macrophages depletion and the *Vibrio* ACD each leave contraction frequency unchanged. We found that *Vibrio’s* T6SS ACD induces host tissue damage and cell death while activating and not specifically killing macrophages. Rather than being selectively killed, intestinal-resident macrophages move from the midgut to the posterior site of tissue damage. These findings point to a parsimonious model: normally, macrophage proximity to enteric neurons downregulates the strength of mechanical contractions via the previously studied BMP2 pathway (21). T6SS/ACD-mediated tissue damage stimulates an innate immune response, recruiting macrophages and thereby removing the downregulatory signal.

From the perspective of the host, this sequence of events provides a simple means of coupling cellular distress to the purging of intestinal contents. From the perspective of *Vibrio*, the purging can displace bacterial competitors leaving it, being motile and planktonic, able to persist in the gut (7, 12, 16). Notably, this connection between bacterial cellular activity and neuromuscular control does not require any special biomolecular signaling pathways, instead harnessing spatial reorganization to reshape intercellular communication. Of course, altered signaling may occur, especially on longer timescales. In *Drosophila melanogaster*, recent work has shown that *Vibrio* can stimulate host BMP signaling in intestinal epithelial cells via its T6SS, and that in zebrafish as well as in fruit flies the T6SS hinders epithelial cell proliferation and tissue repair (32).

Our experiments examine a natural microbial stimulus in a living animal model, implicating dynamic alterations of neuro-immune relationships. Crosstalk between the nervous and immune systems is increasingly realized to be an important aspect of animal function (33–35). In recent years, a variety of chemokines, cytokines, and neuropeptides have been identified as mediators of this crosstalk (33). Though challenging to study, spatial and temporal signatures of inter-system interactions are likely also crucial, given the elaborate geometry of innervation and the motility and morphological plasticity of immune cells. Using intravital imaging in mice, Kulalert *et al*. recently showed that commensal-specific T lymphocyte sphericity, activation, and proximity to nerve fibers are all interlinked, contributing to the skin’s response to its microbiome (36). We suspect that many more spatial aspects of neuro-immune interactions, potentially modulated by host-associated microbes (5), await discovery.

## Acknowledgements

We thank Rose Sockol and the University of Oregon Zebrafish Facility staff for fish husbandry and care, and Adam Fries for imaging advice and assistance. This work was supported by the National Science Foundation under award 2310570 and the National Institutes of Health under award P01GM125576. The funders had no role in study design, data collection and analysis, decision to publish, or preparation.

## Author contributions

All authors contributed to the research design; J.S.N., P.A., J.G.S., and C.T. performed research; K.G., T.J.W., and R.P. supervised research, J.S.N., P. A., and R. P. analyzed data; and J.S.N., K.G., and R.P. wrote the paper.

## Methods

### Animal care

All experiments with zebrafish followed standard procedures (37) and were performed in accordance with protocols approved by the University of Oregon Institutional Animal Care and Use Committee.

### Gnotobiotic Techniques

Wild-type AB, *Tg(mpeg1:mCherry)*, or *Tg(tnfα:GFP);Tg(mpeg1:mCherry)* zebrafish were derived germ-free (GF) and subsequently colonized with bacterial strains, following established protocols (38). In brief, fertilized eggs were collected and incubated in sterile embryo medium (EM) containing 100 μg/mL ampicillin, 10 μg/mL gentamycin, 1 μg/mL tetracycline, 1 μg/mL chloramphenicol, and 250 ng/mL amphotericin B for approximately 6 hours. Following this incubation period, embryos were thoroughly rinsed first in sterile EM containing 0.003% sodium hypochlorite and then in sterile EM containing 0.1% polyvinylpyrrolidone-iodine. These sterilized embryos were then distributed into T25 tissue culture flasks, with a density of one embryo per mL in 15 mL of sterile EM. Notably, during the experiments, the embryos relied on yolk-derived nutrients and were not provided with external feeding. The sterility of the flasks containing larval zebrafish was inspected prior to the experiments.

### Bacterial Growth and Zebrafish Inoculation

*Vibrio* (ZWU0020, PRJNA205585) was previously isolated from the zebrafish intestinal tract. To prepare the bacterial strains for colonization at specific time points, cultures were first grown overnight in Luria Broth (LB) under agitation at a temperature of 30ºC. Then, bacterial cultures were pelleted through centrifugation for a duration of 3 minutes at 7,000 x g, followed by two rounds of washing in sterile EM. An inoculum of 10^6^ colony-forming unit per milliliter (CFU/mL) was used for zebrafish inoculation, directly introduced to the flask water.

### Molecular and Genetic Manipulation

Unless otherwise specified, standard molecular biology techniques were employed. Reagents were used in accordance with the manufacturer’s instructions. We used restriction enzymes and various other reagents for tasks such as polymerase chain reaction (PCR) and nucleic acid modifications. The reagents were primarily obtained from New England BioLabs. To purify plasmids and PCR amplicons, we used kits from Zymo Research. DNA oligonucleotides were synthesized by Integrated DNA Technologies. The sequencing of cloned genes was verified by Sanger sequencing, conducted by Sequetech. A Leica MZ10 F fluorescence stereomicroscope, equipped with a 1x objective lens, was used for the screening of fluorescent bacterial colonies. Genome and gene sequences were retrieved from “The Integrated Microbial Genomes & Microbiome Samples” (IMG/M) website (39).

### Construction of *Vibrio* ZWU0020 T6SS Mutant Variants

The construction of *Vibrio* ZWU0020 mutants containing markerless deletions of either the large T6SS gene cluster or an in-frame deletion of the actin crosslinking domain (ACD) of VgrG-1 was accomplished using allelic exchange and the pAX2 allelic exchange vector as previously described (40). Allelic exchange cassettes targeting each locus were created using splice by overlap extension (SOE). For the large cluster deletion mutant, the following primer pairs were used to amplify 5’ and 3’ flanking homology regions: (5’ HR) 5’-AGTGAAGCAATCGGGCAGT-3’ + 5’-CACACAAAAACATGACTCTGGAAATTTAATTCTGTCCTCATCGGTTAGTAAAATG-3’ and (3’ HR) 5’-CATTTTACTAACCGATGAGGACAGAATTAAATTTCCAGAGTCATGTTTTTGTGTG-3’ + 5’-GGTTCGGTATAGTGGCAGCA-3’. For the ACD deletion mutant, the following primer pairs were used: (5’ HR) 5’-CCCAATGATAGCCACGGTTG-3’ + 5’-CCATTCCATTTTCCACTAGGCTAAAGGACACACCTT-3’ and (3’ HR) 5’-AAGGTGTGTCCTTTAGCCTAGTGGAAAATGGAATGG-3’ + 5’-GGCGCAAGATTTTCAATCA-3’. Each of the resulting 5’/3’HR amplicon pairs were spliced together by SOE and ligated into the pAX2 allelic exchange vector, resulting in pAX2-ZWU0020-T6SSlc (pTW454) and pAX2-ZWU0020-ACD..

Each pAX2 vector was introduced into *Vibrio* ZWU0020 via conjugation utilizing *E. coli* SM10 as a plasmid donor strain as previously described (40). In brief, *Vibrio* and SM10 donor strain were mixed 1:1 on a filter disk and placed on tryptic soy agar (TSA). The mating mixture was then incubated at 30ºC overnight after which bacteria were recovered and spread onto TSA containing gentamicin. These plates were incubated overnight at 37ºC to select for *Vibrio* merodiploids. Isolated merodiploid colonies were screened for successful deletion of the large T6SS gene luster or ACD. Putative mutants were genotyped through PCR with primers flanking each locus to produce differently sized amplicons, representing wild-type and mutant alleles. Primers used for genotyping the large cluster deletion mutant were: 5’-AGTGAAGCAATCGGGCAGT-3’ + 5’-GGTTCGGTATAGTGGCAGCA-3’. Primers used for genotyping the ACD deletion mutant were: 5’-CGGAGCTTTGGTCAATCTCA-3’ + 5’-AGGTCTCTCCGTGGAAAACA-3’.

### Quantifying T6SS-mediated Bacterial Killing in Vitro

Overnight cultures of *Aeromonas* ZOR0001 *attTn7::sfGFP* (40) and either wild-type *Vibrio, Vibrio*^ΔACD^, or *Vibrio*^ΔT6SS^ were mixed at a ratio of 1:3 (*Aeromonas*:*Vibrio*). Mixtures were washed once in 0.7% saline, serially diluted, and spotted onto tryptic soy agar (TSA) (5 ul/spot). Spot dilutions were incubated overnight at 30ºC and imaged with a Leica MZ10F fluorescence stereomicroscope equipped with a K5 Microscope Camera.

### Culture-Based Quantification of Intestinal Bacterial Populations

Dissection of larval guts was done in accordance with established procedures (41). Dissected guts were carefully harvested and transferred to 1.5 mL tubes, each containing sterile 0.7% saline solution and approximately 100 μL of 0.5 mm zirconium oxide beads. A bullet blender tissue homogenizer was used to facilitate homogenization, operated for a duration of 30 seconds at power level 4.

Following homogenization, lysates were serially plated on LB agar plates and incubated overnight at a temperature of 30ºC for enumeration of colony forming units (CFUs) to determine gut bacterial load as previously described (7, 12).

### Measuring Intestinal Motility

Intestinal motility in the form of propagating contractile waves was assessed using Differential Interference Contrast (DIC) microscopy and image velocimetry as previously described (42). DIC videos were recorded at 5 frames per second for five minutes at the distal end of the intestine, indicated in Figure 1A. For analysis, we used open-source particle image velocimetry (PIV) software (43) to calculate a frame-to-frame velocity vector field from the image time-series and then characterized the frequency and amplitudes of gut motions along the anterior-posterior (AP) axis using custom and publicly available software previously described (20) (https://github.com/rplab/Ganz-Baker-Image-Velocimetry-Analysis). We focus on the AP component of the vector field, averaging it along the dorsal-ventral direction to obtain a scalar velocity measure at each position along the gut axis and each time point. The frequency of gut contractions was identified as the location of the first peak in the temporal autocorrelation of motility, while the amplitude of contractions was calculated as the square root of the spatially averaged velocity power spectrum at this frequency, providing a quantitative measure of the magnitude of periodic gut motion.

### Light Sheet Fluorescence Microscopy

Light sheet fluorescence microscopy was performed using a custom-built microscope described in detail elsewhere (44, 45). In brief, a galvanometer mirror rapidly scans one of two laser beams (448 nm and 561 nm, Coherent Sapphire, operating at 5 mW), which is then demagnified to create a thin sheet of excitation light that intersects the specimen. A 40x 1.0NA objective lens positioned perpendicular to this sheet captures the fluorescence emission from the optical section. The sample is then scanned along the detection axis, enabling the generation of a three-dimensional image. To image the entire extent of the intestine, which measures approximately 1200x300x150 microns, we sequentially image four sub-regions and computationally register the images after acquisition. Unless otherwise specified in the text, all exposure times are 33 ms with an excitation laser power of 5 mW. For all light sheet imaging, a 5.5 megapixel sCMOS camera was used. For time series imaging, scans occurred at 20-minute intervals for 13 hour durations.

### Sample Handling and Mounting for Imaging Experiments

Sample mounting followed previously established protocols (16, 44). Larval 6dpf zebrafish were carefully removed from the culture flask and immersed in sterile EM containing 120 μg/mL tricaine methanesulfonate (MS-222) anesthetic. Each specimen was then briefly immersed in 0.7-1.0% low melt agarose drawn into a glass capillary, which was then mounted onto a sample holder. The agar-embedded specimens were partially extruded from the capillary to ensure that the excitation and emission optical paths did not cross glass interfaces. Larvae in the set gel were extruded from the end of the capillary and oriented such that the illumination laser sheet enters from the ventral side. The specimen holder can hold up to six samples simultaneously, all of which are immersed in sterile EM maintained at a temperature of 28ºC. All long-term imaging experiments were conducted overnight, typically beginning in the late afternoon.

### Mounting for Stereoscope Imaging

Each anesthetized specimen was embedded in 4% methylcellulose on a microscope slide. Two stereomicroscopes were used, a Leica MZ10 F fluorescence stereomicroscope equipped with 1x, 1.6x and 2.0x objective lenses or a Nikon SMZ25 stereomicroscope with a 1.0x objective lens.

### *Irf8* sgRNA Cas9 injections

All sgRNAs used were previously designed and characterized with a 100% success rate (46). sgRNAs used are as follows: 5’-ATAAAGCTGAACCAGCGACATGG3’ and 5’ TGGTGAGCAGTCCATGTCAGTGG-3’. The RNA oligonucleotides were resuspended to 20 μM in nuclease free water and stored at -20ºC until use. For *in vivo* applications, 1 nL of an injection mixture composed of 1μL of each sgRNA, 1μL of Cas9, 1μL of Phenol Red (0.125%) were injected into the yolk at the one-cell stage. Phenol Red was used to screen for injected developing embryos during the GF derivation process.

### TNFα and Macrophage Quantification from Dissected Intestines

*Tg(tnfα:GFP);Tg(mpeg:mCherry)* larval zebrafish were mounted for stereoscope imaging. Their intestines were dissected and imaged using a Leica MZ10 F fluorescence stereomicroscope as described above. The number of GFP positive, mCherry positive, and double positive cells were quantified visually using ImageJ.

### Macrophage Image Analysis

Macrophages were identified in two-dimensional fluorescence stereomicroscope images (e.g. Figure 2AB) using custom code written in MATLAB. First, a threshold of twice the median pixel intensity minus the minimum intensity was applied to distinguish potential cells from background. After a 2 pixel morphological opening, connected above-threshold pixels were identified as objects, and those objects with an effective radius outside the range 3-20 μm were eliminated. Separately, a high-pass filter of Gaussian width 6 μm was applied to the original image, and pixels more than five standard deviations above the median were retained and grouped into objects of connected pixels. The high pass filtering helps distinguish cells from particularly bright and broad autofluorescent intestinal background. Cells were identified as the retained objects from either of the two analysis paths, and the resulting output was visually examined to assess its reasonableness.

Macrophages and acridine orange positive cells were identified from three-dimensional light sheet fluorescence microscopy images using custom code written in Python using the open-source packages NumPy, SciPy, scikit-image, and pandas. First, each 2D slice of the z-stack was down-sampled by a factor of 4 in both the x and y directions to reduce segmentation time. Next, background subtraction was performed by subtracting the median value of each 3D image from each pixel value. The images were then denoised using a local median filter. Objects were then segmented using simple thresholding with a threshold value set to the mean plus 10 times the standard deviation of the pixel intensities. To connect pseudopodia to the main cell body, a series of image operations were performed on the segmented objects, including dilation, skeletonization, and object joining, and the resulting output was visually examined to assess its reasonableness. All analysis was performed using custom code written in Python.

### Acridine Orange Staining in Live Larval Zebrafish

To visualize apoptosis in live larval zebrafish, we used the dye Acridine Orange dye (47). Larval zebrafish were derived germ-free and mono-associated with the appropriate bacterial strain. Larvae at 6 dpf were anesthetized then immersed in sterile EM containing 0.3125 μg/mL Acridine Orange for of 10 minutes in the dark. Following the staining, the larvae were washed in sterile EM three times to remove excess dye.

### Data and Statistical Analysis

Custom code written in Python was used for data analysis and plotting. Specific details about the statistical tests conducted for each dataset can be found in the respective figure legends.

