## Supplemental Figures and Movie Captions for "Bacterial Modulation of Intestinal Motility through Macrophage Redistribution"

#### Supplemental Movie Captions

**Supplemental Movie 1:** Gut contractions of a representative 6 dpf zebrafish inoculated with wild-type *Vibrio*. Arrows from image velocimetry analysis are superimposed on images acquired with differential interference contrast microscopy. For size, the movie has been downsampled in space (2x) and time (5x) from the original, analyzed, image set.

**Supplemental Movie 2:** Gut contractions of a representative 6 dpf zebrafish inoculated with *Vibrio*$^{\Delta ACD}$. Arrows from image velocimetry analysis are superimposed on images acquired with differential interference contrast microscopy; the scale of the arrows is the same as in Movie S1. For size, the movie has been downsampled in space (2x) and time (5x) from the original, analyzed, image set.

### Supplemental Figures

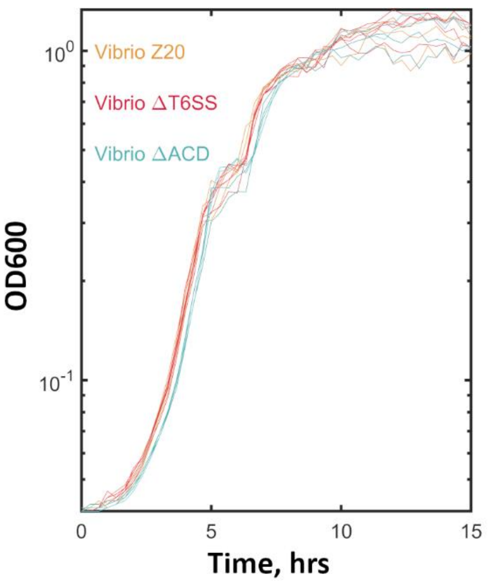


**Figure S1.** Growth curves, measured as optical density in lysogeny broth (LB) at 30 ºC, for wild-type *Vibrio* (*Vibrio* Z20), *Vibrio* lacking the full T6SS gene cluster (*Vibrio*$^{\Delta T6SS}$), and *Vibrio* lacking the actin crosslinking domain (ACD) of the T6SS spike protein VgrG-1 (*Vibrio*$^{\Delta ACD}$), showing indistinguishable growth rates. $N = 4$ replicates were evaluated for each bacterial strain, with optical density measurements fit to logistic growth curves giving growth rates of 0.75 ± 0.02 hr.$^{-1}$ for wild-type *Vibrio*, 0.72 ± 0.03 hr.$^{-1}$ for *Vibrio*$^{\Delta T6SS}$, and 0.74 ± 0.02 hr.$^{-1}$ for *Vibrio*$^{\Delta ACD}$.

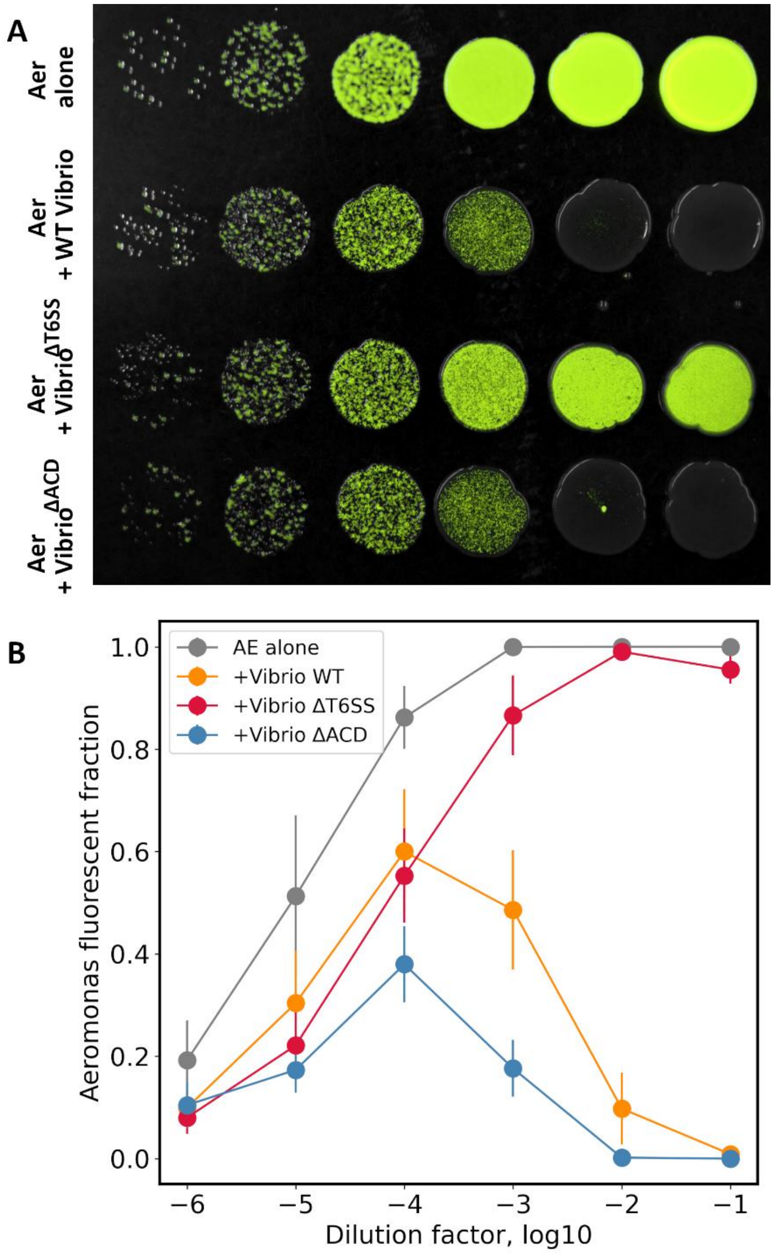


**Figure S2.** In vitro assays of T6SS-mediated inter-bacterial killing. (A) Images of GFP-expressing *Aeromonas* ZOR0001 spotted alone onto tryptic soy agar (top row), or together with wild-type or mutant *Vibrio* strains (lower three rows), at a series of tenfold-increasing initial bacterial concentrations (left to right). (B) Quantification of *Aeromonas* abundance as the fraction of the spotted-disk area showing GFP fluorescence. The mean (solid symbols) and standard error of the mean (error bars) across $N$=6 replicates are plotted as a function of dilution factor. The images and the fluorescent fractions clearly show low *Aeromonas* abundance when *Aeromonas* is co-spotted with either wild-type *Vibrio* or *Vibrio*$^{\Delta ACD}$ and high abundance with *Vibrio*$^{\Delta T6SS}$, consistent with the expectation that deletion of the actin crosslinking domain of the T6SS does not inhibit inter-bacterial killing.

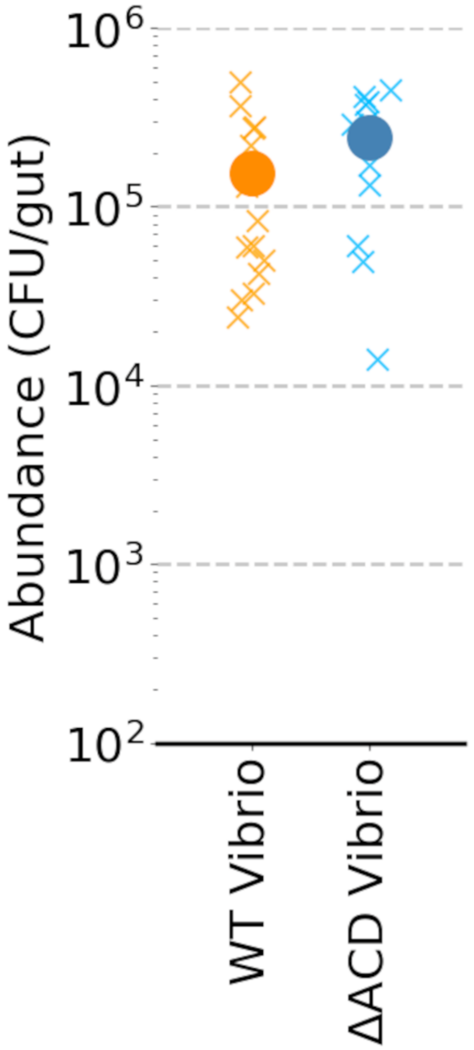


**Figure S3.** Bacterial abundance in the zebrafish gut for *Vibrio*$^{\Delta ACD}$ and the wild-type *Vibrio* inoculated in mono-association with initially germ-free larval zebrafish. Each "x" is derived from plating the dissected gut of a single 6 dpf zebrafish, 24 hours post-inoculation. Solid symbols and error bars indicate the mean and standard error of the mean, respectively. The mean ± standard deviation of $\log_{10}$(bacteria per gut) are 5.0 ± 0.4 and 5.2 ± 0.4 for wild-type *Vibrio* and *Vibrio*$^{\Delta ACD}$, respectively.
